# Massive proliferation of retrotransposons contributes to genome size expansion in species of the *Pseudocercospora* genus

**DOI:** 10.1101/2025.08.13.670038

**Authors:** Sandra-Milena González Sáyer, Ibonne A Garcia, Cristian A Traslaviña, Alex Z Zaccaron, Ioannis Stergiopoulos, Fabio A Aristizabal, Ursula Oggenfuss, Daniel Croll

## Abstract

Genome size expansions are common among eukaryotic lineages. Enlarged genomes can be bioenergetically demanding, and active mobile elements can trigger chromosomal rearrangements and loss of gene function. What triggers genome size expansions remains largely unexplored in many biological clades, particularly within the fungal kingdom. Activation of large transposable elements (TEs), such as long-terminal repeats (LTRs), is a common contributor. Yet the mechanisms of LTR activation remain poorly understood. Here, we focus on the fungal genus *Pseudocercospora* and closely related species with known variation in genome size. In using an assembly-free approach, we found that TE content is highly variable among species, with species-specific retrotransposon families being the main drivers of independent genome expansions. We further focussed on the two species with the most expanded genomes and reference-quality genomes, *P. fijiensis* and *P. ulei*. We found that the *P. ulei* genome is compartmentalized, with highly variable TE densities among chromosomal regions, and a striking reduction in pathogenicity-associated genes. Overall, our study indicates that species of *Pseudocercospora* originally had reduced genome sizes, and genome expansions are species-specific, driven by heterogeneous sets of TE families. Furthermore, we found that in the species with the most expanded genome, TE activity might not have ceased yet as indicated by resequencing data analysis of six strains from diverse locations in Colombia. We discuss what might have caused TE activation and subsequent proliferation in the genus, including stress conditions and host adaptation. Surveys of clades with highly dynamic genome sizes are crucial for the investigation of causal factors driving long-term TE dynamics.

## INTRODUCTION

Eukaryotic species have very diverse genome sizes, covering >10,000-fold changes for haploid genome sizes (1,2). Genome size expansions occurred widely among eukaryotic lineages and may be linked to speciation (3,4). Genome size expansions mostly arise from duplication of existing sequences and are thought to have minimal impact on functional complexity, including the number of gene families or regulatory sequences (5–7). Duplication of regions range in size from whole genome duplications to aneuploidy, to large structural variation and the duplication of genes or transposable elements (TEs) (1,8–12). TEs are mobile genetic units with the ability to translocate or create copies of themselves and subsequently insert into different genomic regions (13,14). TEs are not monophyletic in their origin, and either transpose via RNA intermediates, coding their own reverse transcriptase and creating a copy, or via excision and insertion cycles (15,16). TEs are generally autonomous, and contain all genes needed for their own transposition. However, non-autonomous TEs can transpose by parasitizing autonomous TEs (17). TEs are relatively small, ranging from a few dozen base pairs (bp) in certain non-autonomous elements to several tens of kilobases (kb) for typical retrotransposons, to 700 kb for more complex and clustered TE constructs like Starships (15,18). Individual TE insertions do not increase genome size substantially. However, their ongoing proliferation led, for example, the ∼300 bp *Alu* element to cover almost half of the human genome, small miniature inverted TEs (MITE) proliferation to be responsible for remarkable genome size differences among rice genomes, and retrotransposons to make up around 60 % of the genome of the fungus *Cenococcum geophilum* (19–21). Different stress conditions can be a factor to induce bursts of TE activity, leading to increased copy numbers over short evolutionary timescales. Furthermore, even silenced or non-functional TEs can impact genome size evolution, by inducing large-scale chromosomal rearrangements, duplications or deletions via ectopic recombination (22–24).

At the species and population level, TEs are a mutational force, and the impact of most new TE insertions is deleterious. Consequently, new TE insertions are selected against, and diverse mechanisms exist to suppress TE activity. Eukaryotes use epigenic silencing to prevent TEs from becoming active (25). Epigenetic silencing includes DNA methylation and histone modifications (26,27). Silencing can be reversible, especially under stress conditions (28). In addition, many ascomycete fungi have a defense mechanism against repeats called repeat-induced point mutation (RIP), which induces increased mutation rates in repeats (29). However, individual TE insertion can also have beneficial impacts. For instance, in the wheat pathogen *Zymoseptoria tritici*, a TE insertion upstream of the *Zmr1* promoter regulated the diversity of melanin accumulation (30). TE-derived chromosomal rearrangements in industrial and sea floor adapted strains of *Penicillium chrysogenum* likely increased penicillin production (31,32). Over longer evolutionary time frames, TE derived genes can also become co-opted by integrating into host gene functions (33,34).

The fungal genus *Pseudocercospora* contains over 300 species (35) predominantly consisting of host-specific fungal pathogens that pose a threat to agricultural and natural ecosystems (36). *Pseudocercospora* spp. are globally distributed, with a focus on tropical and subtropical environments (37). Key pathogens include *P. fijiensis*, *P. musae*, and *P. eumusae*, which collectively contribute to the black Sigatoka complex affecting banana crops, and *P. ulei*, responsible for South American Leaf Blight in natural rubber plants *Hevea brasiliensis* (38–40). Despite their significance in agriculture and the environment, only a few *Pseudocercospora* species have been sequenced. The existing genome assemblies strongly indicate that the genus has undergone significant genome size changes (40–45). *Pseudocercospora fijiensis* and *P. ulei* exhibited genomes with sizes of 74 Mb and 93.8 Mb, respectively, which are among the largest genomes in (45) the Capnodiales (21). Both genomes were reported to harbor a significant amount of repeats including TEs (40,44,46). A TE hAT element captured H3 proteins and created 784 copies ultimately targeted by RIP throughout the repetitive regions in *P. fijiensis* (47). Compared to *P. fijiensis*, in *P. ulei* lacks detailed analyses of the TE families responsible of the genome expansion.

Here, we analyzed genome size expansion in the *Pseudocercospora* genus and closely related sister clades. We found *P. musae*, *P. fijiensis* and *P. ulei* to have the most expanded genome sizes, independent of the number of annotated genes. To detect genes potentially involved in the interaction with the host, we made extensive analyses on pathogenicity-associated genes and found a strong reduction in *P. ulei*. We then compared TE content among species and found substantial variation in TE content, indicating ongoing TE activity after speciation. Specific retrotransposons were the primary driver of genome size expansion with distinct contributions to independent genome size expansions. Finally, we found that at least some TE activity persists within *P. ulei* by analyzing genome sequencing data of six strains collected across Colombia.

## RESULTS

### *Pseudocercospora* includes the largest known genomes in the Mycosphaerellaceae

To provide a comprehensive characterization of *Pseudocercospora* spp. genome evolution, we performed a comparative analysis including twenty fungal phytopathogen species grouped within the Mycosphaerellaceae family for which genome assembly and gene annotation data were available in public databases (Supplementary Table S1). Of these, ten species belong to *Pseudocercospora*, i.e. *P. fijiensis*, *P. musae*, and *P. eumusae*, which are part of the Sigatoka disease complex affecting banana crops (41), *P. ulei*, a major threat to natural rubber crops (44), *P. macadamia*, the causal agent of husk spot in macadamia crops (42), *P. cruenta*, responsible for Cercospora leaf spot in cowpea crops, *P. pini-densiflorae*, a cosmopolitan pathogen affecting various pine species, *P. fuligena*, a tomato pathogen causing black leaf mold (43), *P. vitis*, the causal agent of isariopsis leaf spot in *Vitis* spp. crops (48), and *P. crystallina*, a fungal pathogen of eucalyptus crops (35,49). Another eight species from closely related sister clades were included, i.e. *Cercospora beticola*, *C. berteroae*, *C. zeina*, *Sphaerulina musiva*, *Dothistroma septosporum*, *Zasmidium cellare*, *Ramularia collo-cygni* and *Zymoseptoria tritici.* Genome assemblies and annotations of two additional species closely related to *Pseudocercospora*, *Paracercospora egenula* and *Rhachisphaerella mozambica*, were performed in this study.

BUSCO scores for *Pseudocercospora* and sister clade genomes indicated high completeness ranging from 93.8% in *P. musae* to 99.0% in *P. cruenta*, *C. berteroae* and *S. musiva* (Supplementary Figure S1). Genome assembly contiguity (*i.e.*, N50) ranged between 42.9 kb in *P. eumusae* to 5.9 Mb in *P. fijiensis* (Figure 1; Supplementary Table S1). The largest genomes were predominantly represented by the *Pseudocercospora* genus (Figure 1, highlighted in mauve). Genome sizes of all analyzed species ranged from 29.3 Mb in *S. musiva* to 93.7 Mb in *P. ulei*. To compare the number of predicted coding regions per genome, we performed gene annotations for the following eight species to fill gaps in publicly available data, i.e. *P. crystallina*, *P. cruenta*, *P. fuligena*, *P. vitis*, *P. pini-densiflorae*, and the outgroups *Pa. egenula* and *Rh. Mozambica* (annotations available on Zenodo: https://zenodo.org/records/15862053). The number of annotated protein-coding genes in Mycosphaerellaceae species ranged from 7,342 to 16,015 (Figure 1). *Za. cellare* and *P. macadamiae* exhibited the highest counts of annotated genes, with 16,015 and 15,430 genes, respectively. Conversely, *P. vitis* and *P. crystallina* displayed the lowest counts of annotated genes, with 7,342 and 8,716 predicted genes, respectively (Figure 1). The limited number of gene candidates may be attributed to the lack of specific training in the gene prediction algorithm. *Pseudocercospora ulei* has undergone a substantial genome expansion compared to all species in this study. The genome of *P. ulei* was three times larger than its closest relative, *P. vitis*, and 1.2 times larger than *P. fijiensis*, the second-largest genome in this study. The three largest genomes, *P. ulei*, *P. fijiensis* and *P. musae* each showed genome size increase, yet no increase in gene numbers.

**Figure 1.**
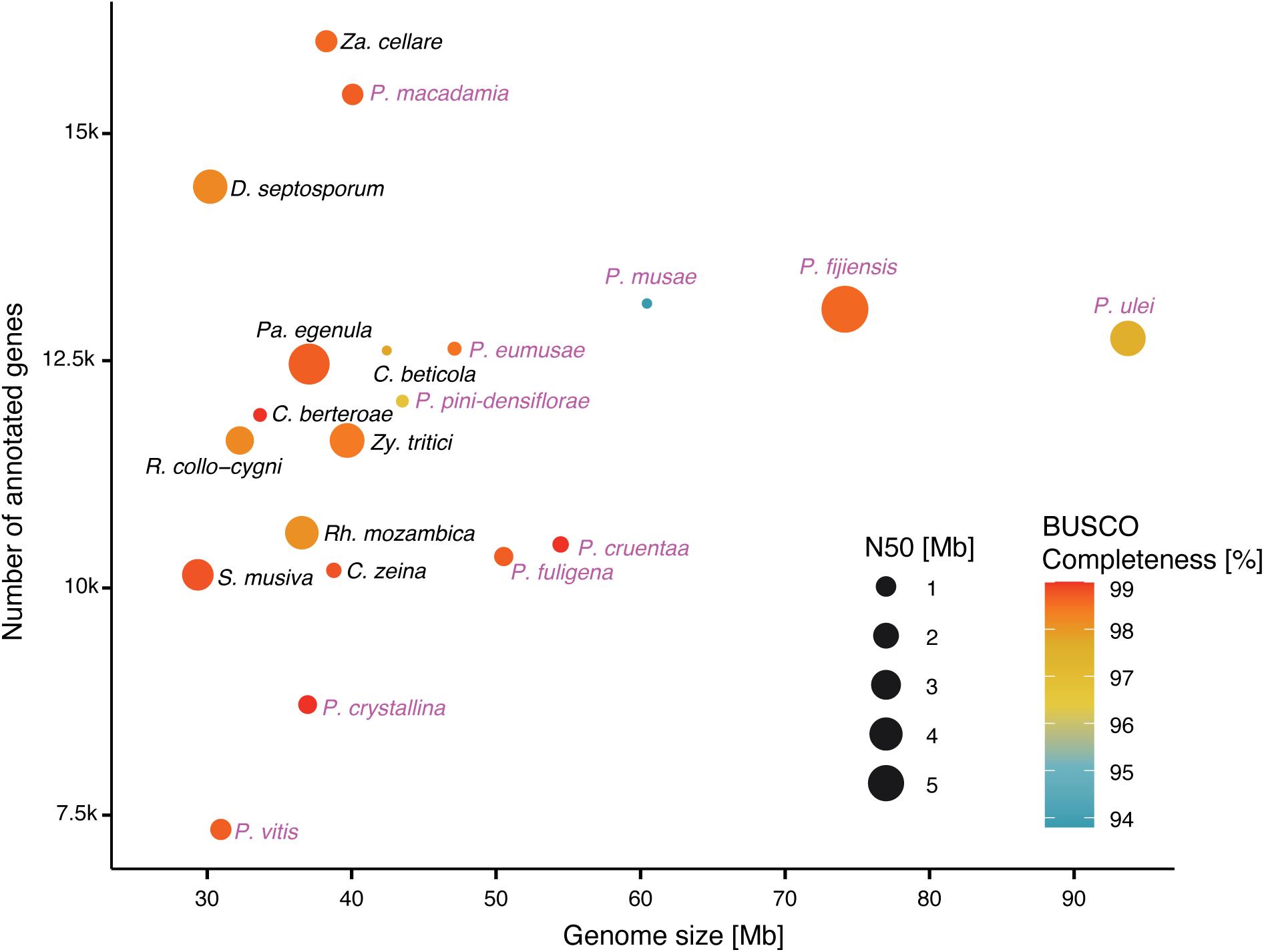
Genome assembly quality metrics of *Pseudocercospora* and closely related species including assembled genome size, gene content and N50 values. The dot color represents to genome assembly completeness score, which was assessed with the number of complete single copy BUSCO (Benchmarking Universal Single-Copy Orthologs) genes. Dot size indicates N50. Genomes from the *Pseudocercospora* genus are labeled in mauve.

### Phylogenomic analyses reveal independent genome size expansions

The phylogenetic relationship of the *Pseudocercospora* genus and closely related species within the Mycosphaerellaceae was assessed using *Blumeria graminis* from the Erysiphaceae family as an outgroup to root the tree. The species grouped into three distinct clades as expected (Figure 2A). Clade A contained most of the *Pseudocercospora* species, except for *P. crystallina*. The two newly assembled species *Rh. mozambica* and *Pa. egenula* clustered with clade C. The species closest to massively expanded *P. fijiensis* and *P. ulei* genomes each showed small genome sizes.

**Figure 2.**
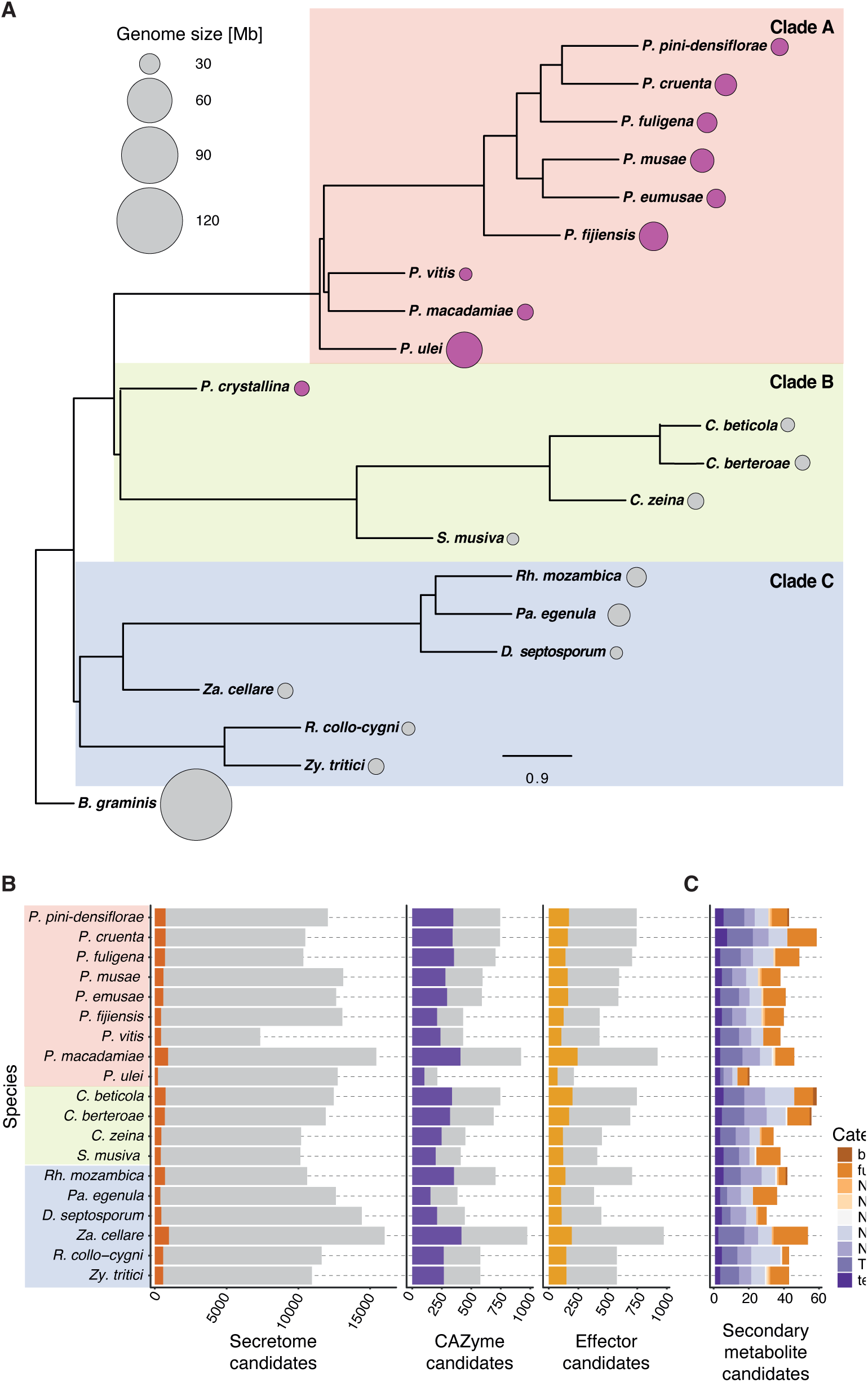
Phylogenetic relationship and pathogenicity-associated genes in Mycosphaerellaceae family genomes. A) Phylogenetic tree of species within the Mycosphaerellaceae family. Dot plots represent the genome size. Genomes belonging to the *Pseudocercospora* genus are filled in mauve. *Blumeria graminis* was used to root the tree as an outgroup. B) Secreted protein profiles of species within the Mycosphaerellaceae family. Left: The gray background represents the total proteome, and dark orange indicates the predicted secretome. Middle and right: The gray background represents the predicted secretome, while carbohydrate-active enzymes (CAZyme) are shown in purple and effector candidates are shown in yellow. C) Secondary metabolite gene clusters in species of the Mycosphaerellaceae family. The colors indicate the different categories of secondary metabolite gene clusters.

### Reduction in pathogenicity-associated genes in *P. ulei*

To assess whether genomes experienced an expansion of pathogenicity-associated genes, we estimated the number of candidates for the secretome, with a focus on carbohydrate-active enzymes (CAZymes) and effectors (Figure 2B). As expected, the secretome candidates made up only a small share of the entire proteome. We identified differences in the *Pseudocercospora* genus with the largest secretome in *P. macadamiae* (*n* = 921 proteins) and the smallest in the closely related *P. ulei* (*n* = 212). *Pseudocercospora vitis* presented a reduced proteome with a similar number of secretome candidates compared to the genus. *Pseudocercospora ulei* showed a reduced number of CAZymes (*n* = 102) and effector candidates (*n* = 73) as well. However, CAZymes and effectors made up a larger proportion of the secretome compared to the other *Pseudocercospora* and closely related species. Secondary metabolite gene clusters showed similar numbers and proportions in categories among the genus (Figure 2C). *Pseudocercospora ulei* also showed a reduced number of secondary metabolite gene clusters, with a proportionally strongest reduction in T1PKS and NRPS categories. The number of pathogenicity-associated genes may be correlated with lifestyle in fungi. *Pseudocercospora ulei* was described as a biotroph, while the other species are likely either necrotrophic or hemibiotrophic. Consistent with total gene content, pathogenicity-associated genes and gene clusters were not correlated with genome size expansions in *P. fijiensis* and *P. ulei*.

### Genome size increases associated with TE expansions

To objectively compare the repeat content among *Pseudocercospora* and closely related species, we used an assembly-free approach based on short read sequencing. Such an approach likely underrepresents repeat content but removes the bias stemming from unequal genome assembly qualities. We used the tool dnaPipeTE for assembly-free repeat detection assessing the following repeat types: low complexity, rRNA repeats, simple repeats, and TEs. TEs were further classified into LTR retrotransposons, LINEs and DNA transposons. The repeat content varied strongly within the *Pseudocercospora* genus and closely related species, ranging from 1.63% in *P. macadamiae* to 71.02% in the closest relative *P. ulei* (Figure 3, Supplementary Figure S2A). *Pseudocercospora macadamiae*, *P. pini-densiflorae* and *Rh. mozambica* showed very low repeat contents of less than 5%, all of which have small genomes. In *P. cruenta* and *P. eumusae* around a quarter of the genome was covered by repeats, and in *P. musae, P. fijiensis* and *Pa. egenula*, around half of the genome was covered by repeats. Generally, closely related species showed drastically different repeat contents. We observed a significant correlation (Pearson’s, *r* = 0.8, *p* = 0.01) between genome size and the proportion of repetitive sequences across *Pseudocercospora* species, indicating that genome size expansion is likely driven largely by the proliferation of repetitive elements. Additionally, we found a strong negative correlation (*r* = −0.84, *p* = 0.004) between the repetitive content and the proportion of pathogenicity-associated genes in *Pseudocercospora* genomes. Most repeats in genomes with moderate to high repeat content were unclassified TEs. Failure for classification by dnaPipeTE likely stems from fragmentation or low coverage. We found variation in TE lengths among genomes with *e.g*., most TEs being below 500 bp in *P. ulei* (Supplementary Figure S2B). Given that LTR retrotransposons can vary between a few kb to tens of kb, the small repeats lengths in *P. ulei* indicates nested TE insertions resulting in fragmentation. LTR retrotransposons remained at low proportions, except for *P. ulei*, where these accounted for ∼30% of the genome. Other TE types were only detected at low proportions. LINEs were only detected in *P. eumusae* and *P. ulei*, and DNA transposons were only detected in *P. musae*. Simple repeats expanded slightly in *P. cruenta* and *P. ulei*. Repeats of low complexity and rRNA remained at low proportions throughout the genus. Neither the repeat content nor the types of repeat content correlated with the phylogenetic position of the species, indicating likely independent bursts of repeat activation.

**Figure 3.**
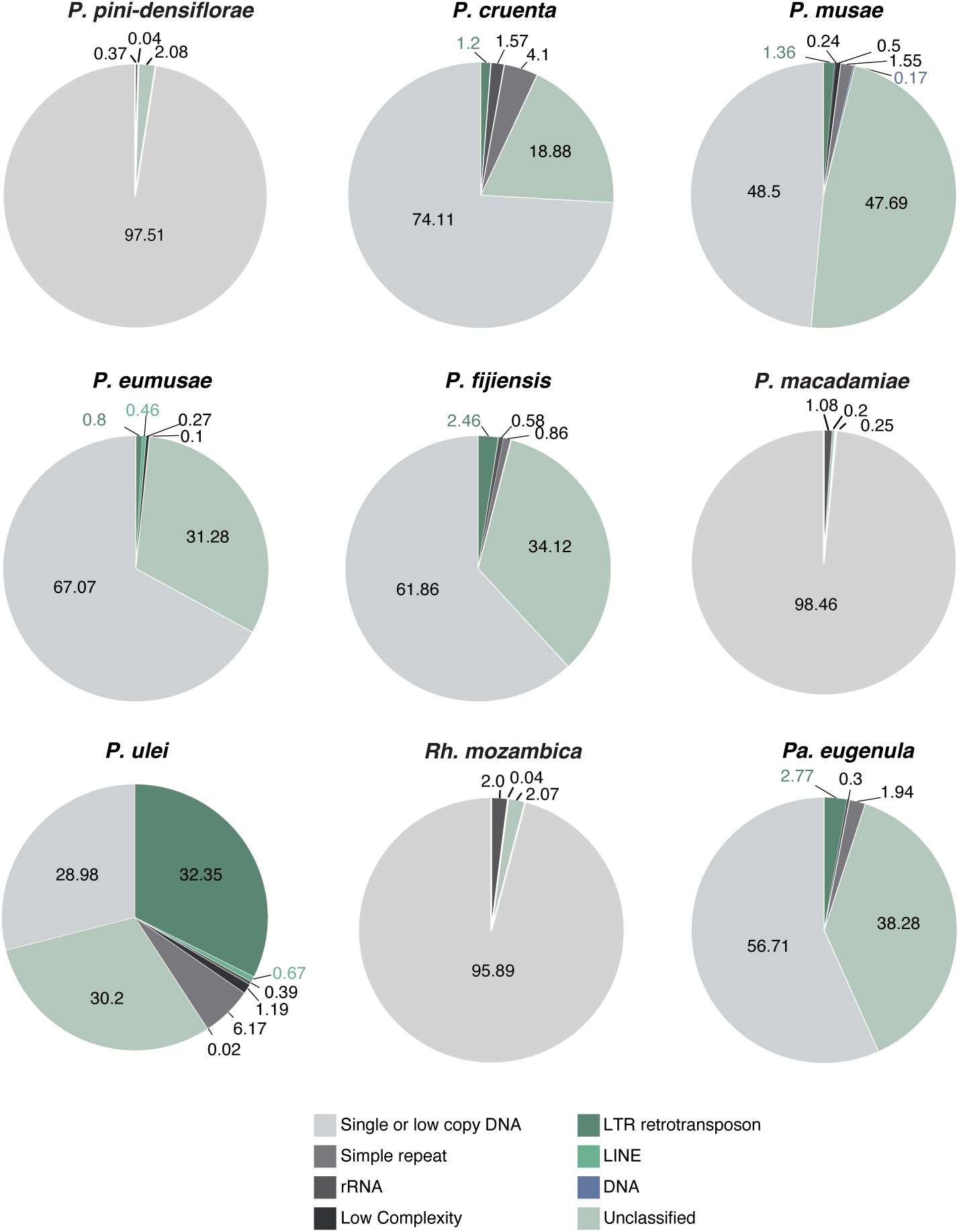
Repeat element distribution among *Pseudocercospora* genus genomes and two closely related species. Light gray indicates the estimated non-repetitive portion of the genomes. Dark grays indicate repeats of low complexity, rRNA and simple repeats. Green and blue colors indicate TEs. The order of the plots follows largely the phylogenetic grouping (Figure 2A).

### LTR retrotransposons underpin genome size expansions in *Pseudocercospora*

The assembly-free TE detection with dnaPipeTE indicates strong and phylogeny-independent TE expansions. To further clarify the genome size expansion dynamics, we used the Earl Grey pipeline to produce high-quality TE annotations and classifications for the two high-quality expanded genomes of *P. fijiensis* and *P. ulei*. TE coverage in the genome assessed by Earl Grey was similar to the estimations by dnaPipeTE (Figure 4A; Earl Grey TE family consensus sequences and annotations are available on Zenodo: https://zenodo.org/records/15862053; TE family names are species specific and do not indicate shared families). However, classification of the individual TE families was improved by access to the full genome sequence. Length estimates of TE fragments were also more robust. Most TE copies belong to LTR retrotransposons consistent with the assembly-free TE detection approach. In *P. fijiensis*, LTR retrotransposons covered 29.4% (21.8 Mb) of the genome, followed by unclassified TEs (13.3%, 9.8 Mb), DNA transposons (3.9%), Rolling-circle/Helitrons (2.7%), and LINE (1.7%). In *P. ulei*, LTR retrotransposons made up the largest part of the genome (74%, 69.5 Mb), followed by a small fraction of unclassified TEs (3.1%, 2.9 Mb), and satellite sequences (0.5%). The bulk of the detected TE fragments overall, which includes full-length elements and fragmented ones due to nested insertions were either LTR retrotransposons or remained unclassified (Figure 4B). The diversity of TE superfamilies is larger in *P. fijiensis* (*n* = 22), while the *P. ulei* genome includes only 8 TE superfamilies. Notably, most LTR retrotransposons belong to the RLG superfamily (formerly known as *Gypsy*, and to be renamed, see (50)), with 4,787 TE fragments in *P. fijiensis* and 26,795 TE fragments in *P. ulei*. Other LTR retrotransposons were only found at low copy numbers. Given the evidence for high degrees of TE fragmentation, we compared the lengths of each LTR retrotransposon fragments between the two species and *Z. tritici*, which was subject to an extensive manual TE curation (51,52). Consistent with the assembly-free approach, the mean length of LTR retrotransposons was lower in *P. ulei* compared to the other species, however, most TE fragments were found to be >500 bp (Figure 4C). Mean LTR retrotransposon length was 3,996 bp in *P. fijiensis*, 2,569 bp in *P. ulei*, and 3,790 in *Z. tritici*.

**Figure 4.**
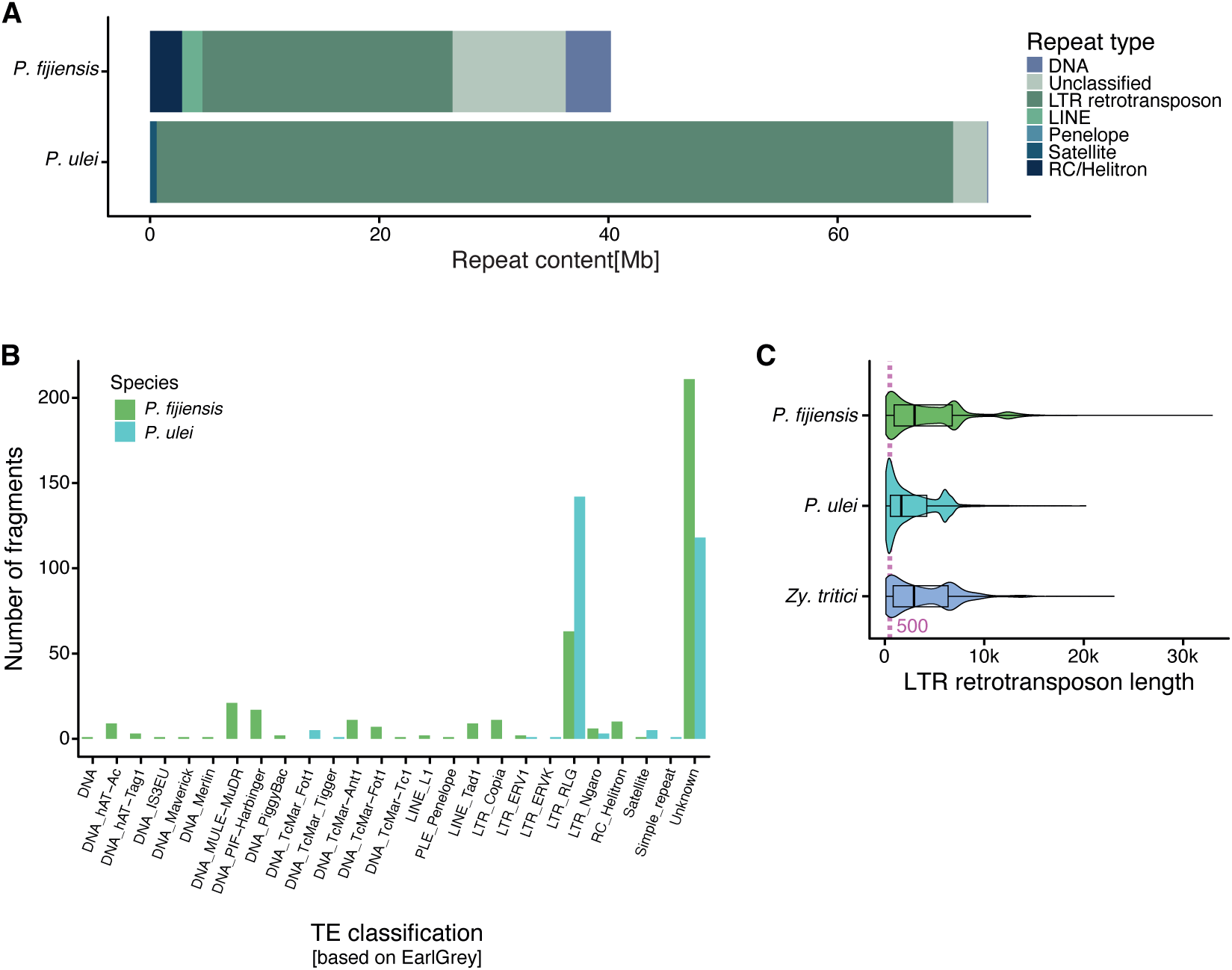
TE coverage in the two enlarged genomes of *P. fijiensis* and *P. ulei* based on high-quality detection and classification with Earl Grey. A) Total length of TEs per genome. The colors indicate the TE category. B) Number of TE copies assessed by Earl Grey for the different TE superfamilies. Copies were counted as both full-length TEs or fragments. Colors indicate the species. C) Distribution of LTR retrotransposon fragment lengths detected in *P. fijiensis* and *P. ulei* compared to the manually curated TE content of the *Z. tritici* outgroup. TEs include both full-length and fragmented TEs. The mauve line indicates 500 bp (as in Supplementary Figure S2B that shows length distribution of any repeat discovered with dnaPipeTE).

The genome-wide analysis of TEs revealed 391 species-specific families for *P. fijiensis*, and 277 for *P. ulei*, with no shared TE families. To identify candidate TE families responsible for recent TE activity bursts and subsequent genome size expansion, we filtered for families with copy numbers above 200. The *P. fijiensis* genome showed only two TE families with copy numbers above 200, but we found 29 such TE families in *P. ulei*, one of which had >800 copies (Figure 5A). The high-copy TE families in *P. fijiensis* belong to RLG and Helitrons, and RLG in *P. ulei*, with two TE families remaining unclassified. Taken together, this suggests that the repeat expansion in *P. fijiensis* likely stems from a higher diversity of low copy TEs, while the high-copy numbers of a few TE families in *P. ulei* suggests a more recent burst of fewer TE families.

**Figure 5.**
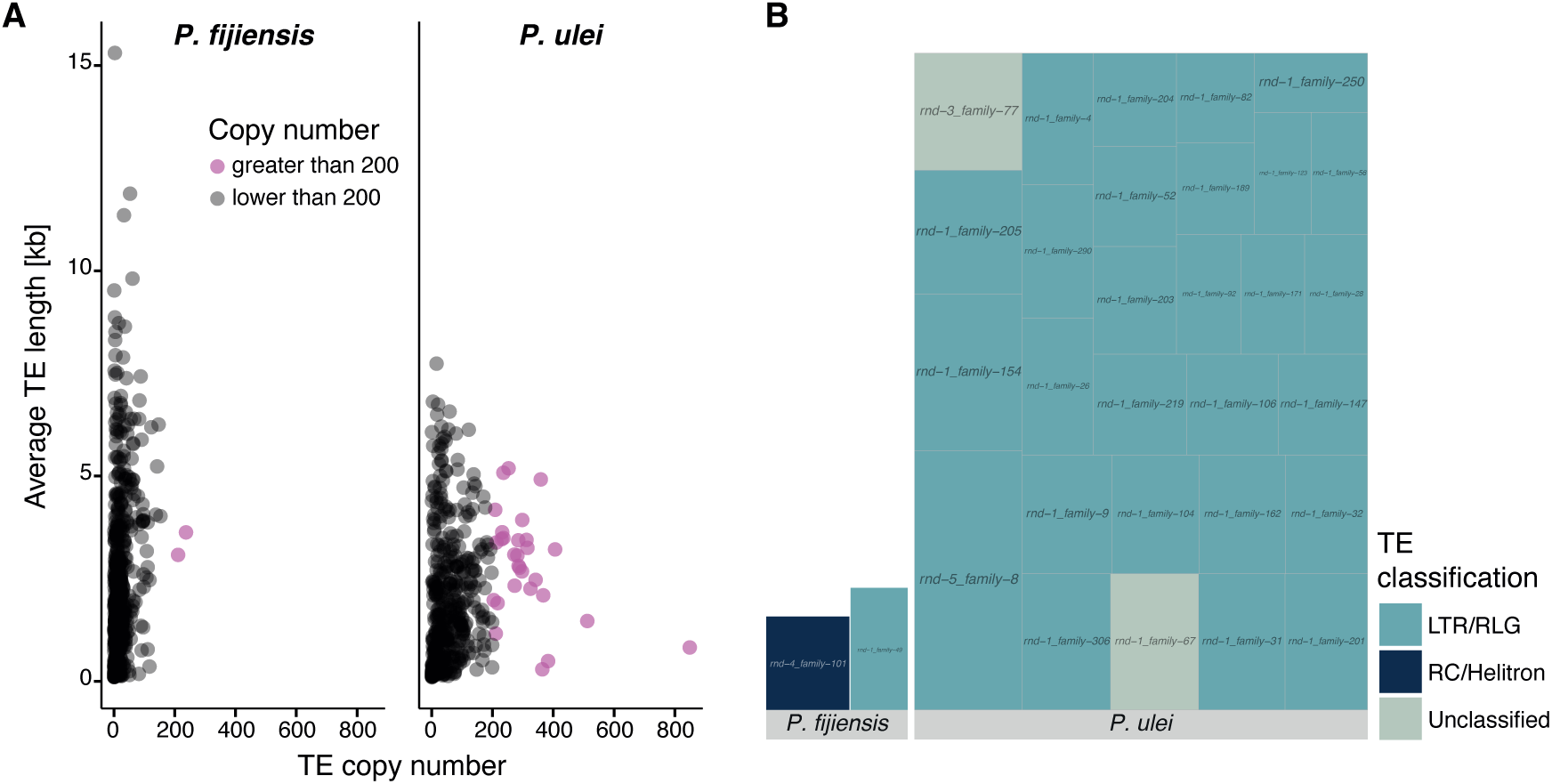
TE families with high copy numbers in *P. fijiensis* and *P. ulei* based on high-quality detection and classification with Earl Grey. A) Correlation of copy number and average length of species-specific LTR retrotransposons. TEs include both full-length TEs and fragments. Mauve dots indicate TE families with more than 200 copies. B) Copies of high-copy number TE families annotated in *P. fijiensis* and *P. ulei*. Colors indicate the superfamily. Each TE family is indicated by a box, and box size indicates the number of copies.

To improve the TE classification and to reduce fragments of TEs erroneously classified as full-length elements, we conducted a manual curation of TE families in *P. ulei*, created consensus sequences and renamed the remaining TE families according to the three-letter code from Wicker et al (2007). Many RLG or unclassified families detected by Earl Grey were mostly fragments of the newly named RLG_Mira family, followed by RLG_Ginan. Given the estimated length for the TE consensus sequences, we confirmed that most TEs in *P. ulei* were fragments of less than 80% full-length (Supplementary Figure S3A). Among high-copy retrotransposons, lengths remained highly variable, especially for the two high-copy TE families RLG_Mira and RLG_Ginan (Supplementary Figure S3B). We created a phylogenetic tree for RLG_Mira coding regions and identified two well differentiated clusters (Supplementary Figure S3C). GC content of RLG_Mira coding regions showed almost exclusively a moderate to high GC content, indicating they the element was not affected by RIP despite being recently active.

### Genomic landscape of the reference-quality *Pseudocercospora* genomes

To identify TEs located close to genes, we calculated the distances between annotated genes and the closest TE (Figure 6A). TEs were generally closer to genes in *P. ulei* (mean = 14,922 bp) compared to *P. fijiensis* (mean = 17,785 bp) and *Z. tritici* (mean = 68,130 bp). Only a small number of direct overlaps were detected in *P. fijiensis* (n = 98, 0.8% of all genes) and *Z. tritici* (n = 31, 0.3% of all genes), however, significant overlaps were found in *P. ulei* (n = 6,441, 51.1% of all genes). Furthermore, we analyzed gene, TE in general and RLG_Mira contents in windows of 10 kb for the largest 12 scaffolds in *P. ulei* (Figure 6D). We found a strong compartmentalization between TE-rich regions with a reduced number of genes and gene-rich, TE-depleted regions. RLG_Mira elements were present in most TE-rich regions, but differed in the amount of overlap. Next, we overlayed large RIP affected regions, which showed a similar distribution as the TE-rich regions. Effector and CAZyme candidates were detected in each of the compartment types. The strong compartmentalization of the *P. ulei* genome indicates strong purifying selection acting against new TE insertions in gene-rich regions, and relaxed selection in TE-rich regions. The proximity of TEs and some genes could also stem from some misannotated genes being in rather TEs.

**Figure 6.**
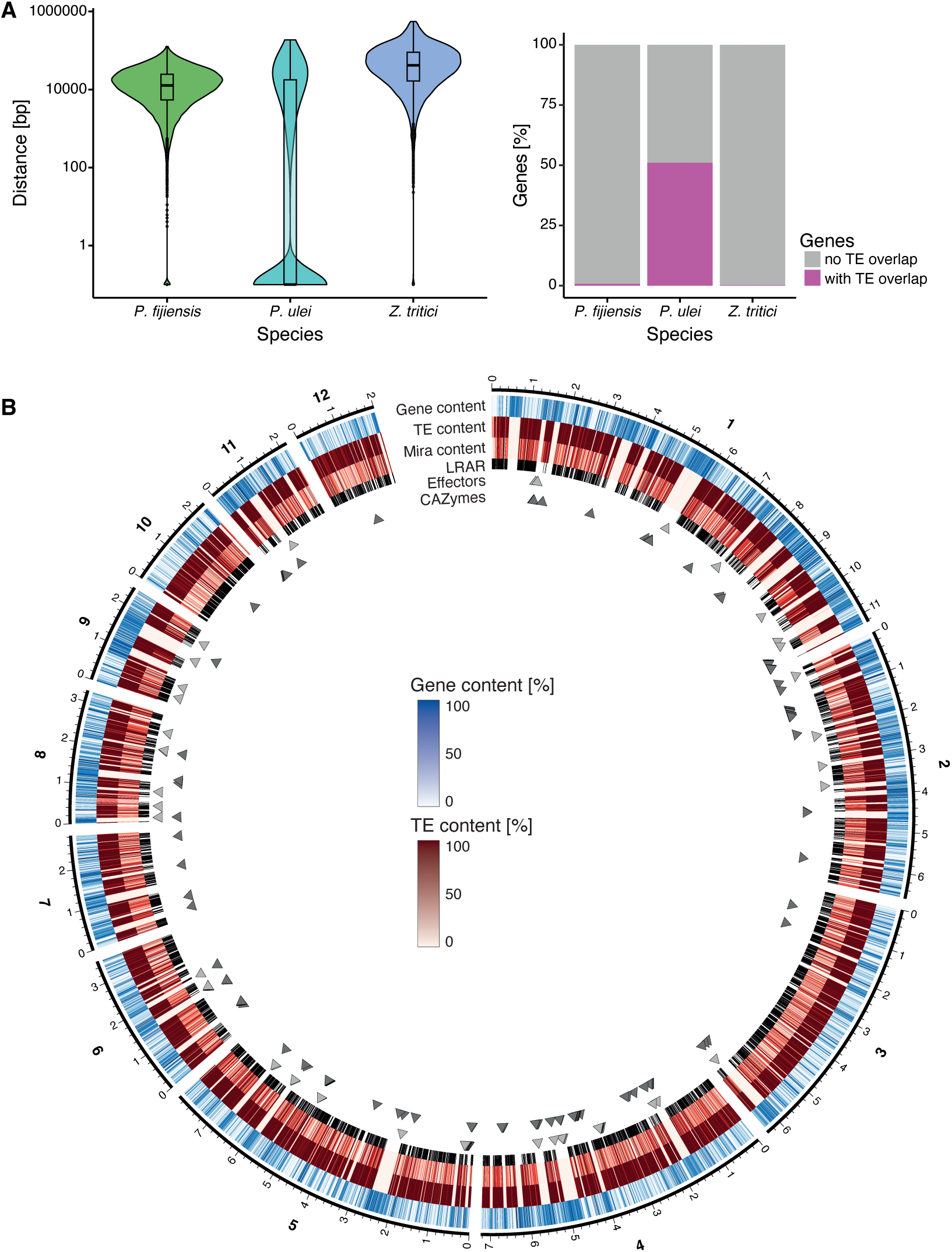
TE landscape in *P. fijiensis*, *P. ule*i and *Z. tritici*. A) Variation in distances between genes and the closest TE. Genes with more than one TE insertion were treated as a single insertion. B) Circos plot visualizing the genomic landscape for *P. ulei*. Genes, TEs and RLG_Mira content were calculated for window sizes of 10 kb. Large RIP affected regions (LRAR) indicate regions of at least 4 kb with a high RIP composite index.

### TE content analyses of *P. ulei* strains

To determine whether some TE activity persist in *P. ulei,* we sampled and whole-genome sequenced strains from natural rubber tree infections in three different locations across Colombia (Figure 7A). We assessed the genetic structure of the *P. ulei* strain collection using 1,802,029 genome-wide SNPs. Strains clustered into three distinct groups according to geography (Figure 7A). We then mapped short-read data against the manually curated *P. ulei* consensus library to assess the coverage and estimate the number of copies for each TE family. TE copy numbers were largely stable and similar to the direct assessment in the reference genome (Figure 7B). Two strains had a slightly higher number of estimated TE copy number than the reference strain. Estimated TE copy numbers varied slightly, but independent of geographic origin. Like the reference genome, retrotransposons and the RLG superfamily were overrepresented in the additional strains as well. At the TE family level, RLG_Mira and RLG_Ginan families were the predominant components of the repetitive content of all the *P. ulei* strains assessed, as seen in the reference genome (Figure 7C). We observed small differences in TE family coverage among the six strains and the *P. ulei* reference genome data, however these differences may reflect limitations in TE detection with short reads rather than biological variation. High TE content is, hence, a broadly shared pattern within the species, and variability in content indicates that TE activity might be ongoing.

**Figure 7.**
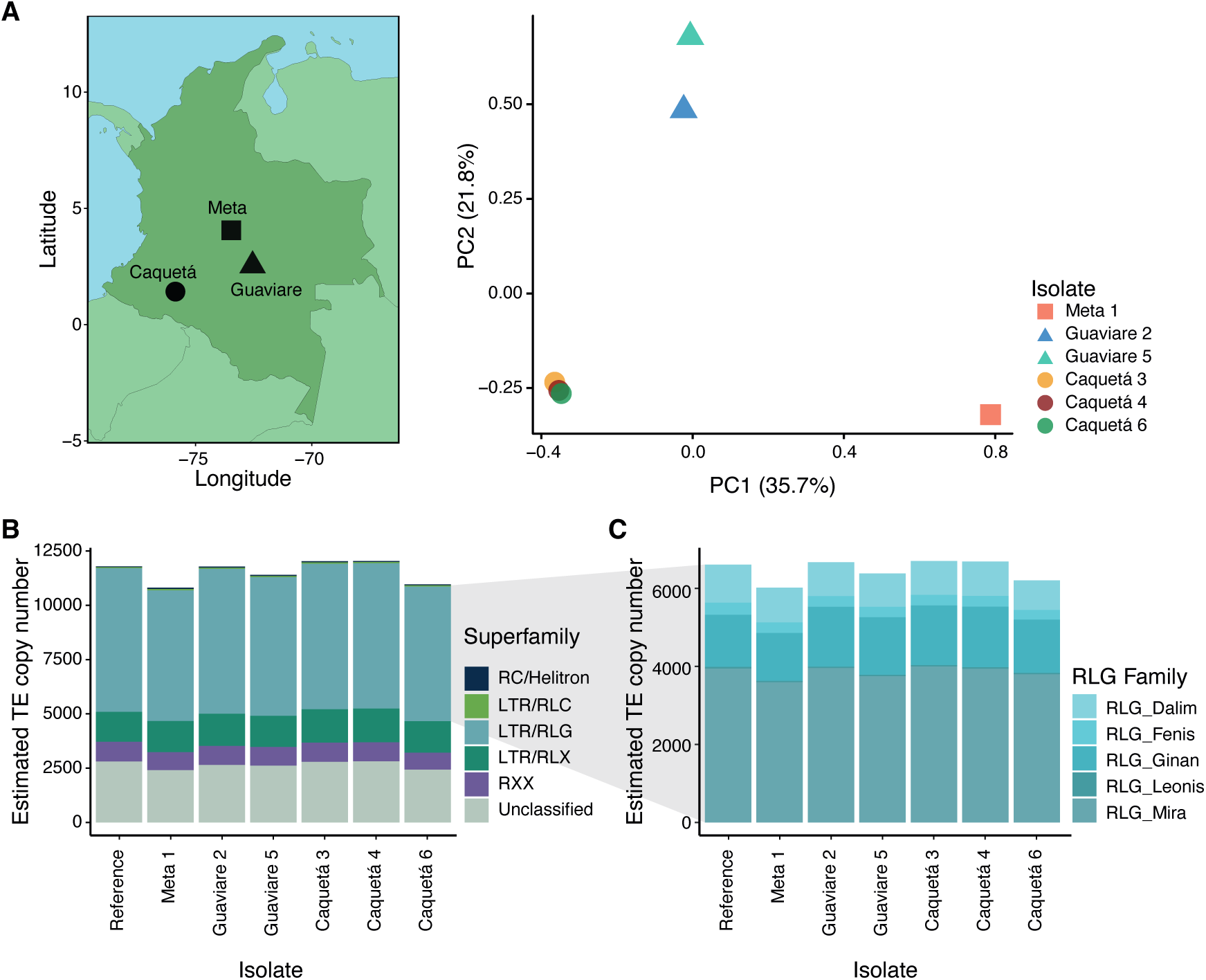
Whole-genome sequencing and TE content analyses of six *P. ulei* strains collected in Colombia. A) PCA analysis of *P. ulei* strains based on linkage-pruned genetic variants and their geographical locations. B) Estimated coverage of *P. ulei* strains using the McClintock coverage analysis based on a curated TEs library for the genus. The color represents the superfamily. C) Estimated coverage of species-specific RLG TE families.

## DISCUSSION

Tracking genome size evolution among fungi remains limited to few groups including the Pucciniales, Erysiphaceae or Glomeraceae (53–57). Our study aimed to explore genome size dynamics within the species-rich *Pseudocercospora* genus consisting predominantly of host-specific plant pathogens (35). We found highly variable genome sizes even among closely related species. Our assembly-free approach based on low coverage short reads allowed us to create draft repeat coverage estimations that could be confirmed with high-quality genome assemblies.

Genome enlargement did not correlate with a higher number of genes or pathogenicity-associated genes. The largest genome, *P. ulei,* even showed a slight reduction in coding sequence content, and a dramatically increased number of TE insertions into predicted genes. Genome expansions were largely caused by TE expansions, but the expansion characteristics varied in terms of number and diversity of TE families involved. RLG retrotransposons were consistently involved in genome size expansions, yet no TE family was involved in more than one observed burst. Genome size expansions are likely phylogeny-independent, and might be caused by the activation of specific TE families during stress conditions, or by horizontal transfer of TEs with subsequent bursts in the new host.

To compare genomes of various quality and annotation, we used an assembly-free approach with short reads. dnaPipeTE detects TEs even in low coverage genomes. The repeat contents of the genomes remain a rough estimation, and most of the potential TE families were not classified. The use of short reads made it harder to estimate the coverage of the genome by TEs and other repeats, as repeats are too short to cover most TEs, and the presence of a TE fragment carries no information about the size of the full-length TE. However, when using the assembly based TE detection with Earl Grey in the two best assembled genomes, i.e. those of *P. fijiensis* and *P. ulei*, we found similar repeat contents. Earl Grey estimated larger numbers of classifiable TE families. However, the short read approach was not useful to fine-tune TE classifications or clearly assess locus specific TE insertions. In contrast, the tool-provided coverage analyses of the genomes provided a less skewed variation in TE content.

Our findings indicate that genome size expansions among *Pseudocercospora* species were largely caused by differences in TE content. We found a striking difference in genome size and repeat content between each *P. ulei* and its closest relative *P. macadamiae*, and between *P. fijiensis* and its closest relatives *P. musae* and *P. eumusae*. In addition, different, species-specific TE families were responsible for these individual expansions. *Pseudocercospora ulei* had an almost exclusive expansion by LTR retrotransposons, namely by the two families RLG_Mira and RLG_Ginan. *Pseudocercospora fijiensis* also had an expansion of RLG elements, although DNA transposons and LINEs were part of the expansion as well, the number of TE families was higher, yet the copy per family was a bit lower. *Pseudocercospora fijiensis* TEs are known to be severely impacted by the defense mechanism RIP (46). Even though *P. ulei* shows many large RIP affected regions, the expanded RLG_Mira coding sequences show no indications of RIP effects. This raised the question whether RIP is still functional in *P. ulei*. Recent losses of the RIP machinery were detected repeatedly among ascomycetes (58).

The TE induced genome size expansions were most likely initiated by independent triggers, resulting from activated TE families not shared by a direct ancestor. TE activity is often caused by stress conditions, in which previously silenced TEs are de-repressed and create new copies (28,59). For fungal plant pathogens, stress includes the response of the host, fungicide application or climatic factors. Such stressors might have induced TE activity, initiating stepwise genome size increases. Expansions would have been countered if TEs were inserted frequently into conserved regions, creating strong negative effects. Fungi might experience elevated stress when adapting to a new host species. *H. brasiliensis*, the host of *P. ulei*, contains a poisonous latex with antifungal properties (60). Depending on which host the last shared ancestor of *P. ulei* and *P. macadamiae* occupied, one or both species might have encountered similar new stress conditions.

The strong clustering of TEs into chromosomal compartments in *P. ulei*, and the large amount of fragmented TEs rather than retention of full-length copies indicates that existing TE-rich regions are preferred insertion sites for new TEs, by this contributing to fragmentation. *Pseudocercospora ulei* shows a reduced set of effectors, a characteristic shared with *Oidium heveae*, another pathogen of *H. brasiliensis* (61). Further examples do not follow this same pattern though with other fungal and oomycete pathogens of *H. brasiliensis* showing no reductions, or even increases in effector gene content. Hence, reduction of effector genes is unlikely to be a general adaptation to the host plant (62–66). Genus-wide analyses and beyond can identify broad patterns of factors influencing TE activity. Our study reinforces the observation that genome size expansions are initiated mostly in terminal branches making direct observation of causal factors very challenging. However, broad surveys of agriculturally relevant plant pathogen genera will build towards a more complete picture of the interplay of TEs, genome sizes and pathogen functions.

## METHODS

### Data acquisition and genome assembly

We obtained genome assemblies for 20 species of the Mycosphaerellaceae family from the National Center for Biotechnology Information (NCBI, Supplementary Table S1). Our comparative genome analysis focused on ten species from public databases from the *Pseudocercospora* genus: *P. cruenta*, *P. crystallina*, *P. eumusae*, *P. fuligena*, *P. fijiensis*, *P. musae*, *P. macadamiae*, *P. ulei*, *P. pini-densiflorae*, and *P. vitis*. An additional eight related species genomes were accessed from public databases: *C. beticola*, *C. berteroae*, *C. zeina*, *S. musiva*, *D. septosporum*, *Za. cellare*, *R. collo-cygni* and *Z. tritici*. Because no assembly was available, we used short-read sequencing data to produce draft genome assemblies for two additional closely related species, *Pa. egenula* and *Rh. mozambica*. *Pa. egenula* and *Rh. mozambica* genome assemblies were constructed from Illumina short-reads. For that end, Illumina raw reads were initially assessed with FastQC v.0.11.9 (https://www.bioinformatics.babraham.ac.uk/projects/fastqc/). Reads were pre-processed with Trimmomatic v.0.39 using the following parameters: ILLUMINACLIP:TruSeq3-PE-2.fa:2:30:10:1:true TRAILING:2 SLIDINGWINDOW:4:15 MINLEN:90 (67). For phylogenetic analyses, we included *B. graminis* as an outlier group to root the tree (68). Trimmed reads were assembled with SPAdes v.3.13.0 using the--careful parameter (69). Assemblies were evaluated with QUAST v.5.2.0 (70). Genome completeness was assessed based on orthologous gene composition obtained with BUSCO v.5.7.1 using the ascomycota_odb10 database (71).

### Phylogenetic analyses

We performed phylogenomic inference using 1315 orthologous BUSCO gene sequences that were shared by all genomes. Amino acid sequences from single copy orthologous genes shared between the assessed species were aligned with MAFFT v.7.310 with the parameters --genafpair -- maxiterate 1000 (72). A maximum likelihood tree was estimated with IQTree v.2.2.5 using 1000 bootstrap replicates (73). For this, we initially concatenated independent tree files previously inferred and then estimated the tree with Astral v.5.7.8 using the default parameters and *B. graminis* as outgroup to root the tree (74).

### Structural and functional annotation

To compare functional annotations among genomes, we performed structural annotations for genomes were this information was lacking: *P. crystallina*, *P. cruenta*, *P. fuligena*, *P. vitis*, *P. pini-densiflorae*, *Pa. egenula* and *Rh. mozambica*. For this, we used Augustus v.3.5.0 (75), training the gene predictors with gene models from the *P. ulei* reference genome (44). Predicted proteomes were extracted from the gene candidate catalogs using gffread v0.12.1 (76). To predict proteins interacting with the plant host, we used the Predector pipeline v.1.2.7 (77). Predector integrates a range of fungal secretome and effector discovery tools, and ranks the effector candidates based on machine learning methods. Predector also includes CAZyme identification which is predicted by homology mapping the amino acid sequences against dbCAN v.10 database (78). Secondary metabolite gene clusters were predicted from genome assemblies using the antiSMASH web server v.7.0 (79).

### Transposable element annotations

Given the variable quality of genome assemblies in the *Pseudocercospora* genus, we resorted to analyzing TEs with dnaPipeTE, an assembly-free tool optimized for TE detection in low coverage short read datasets (80). Briefly, dnaPipeTE analyzes repetitive elements by processing raw genomic reads. It selects three subsamples of low coverage (<1x) reads to assemble TEs into contigs. The assembled contigs are then annotated through homology comparisons using the DFAM database (81). DnaPipeTE was used for TE annotation only in the eight *Pseudocercospora* species and two closely related species *Pa. egenula* and *Rh. mozambica* for which short-read sequencing data were available (Supplementary Table S1).

### Sampling of *P. ulei* strains

We gathered a total of six *P. ulei* strains isolated in the main rubber producer regions of Colombia (Figure 7A). *Pseudocercospora ulei* strains were isolated from *H. brasiliensis* clones established in different regions of Colombia. Two strains were isolated from the leaves of the FX 3864 clone located in Vereda Santa Rosa in the Guaviare Department and provided by the Guaviare Rubber Producers and Marketers Association. Three strains were isolated from leaves of the IAN 873 clone located in the municipality of Belén de los Andaquíes on the Los Gomas farm and provided by the Universidad de la Amazonia. One strain was isolated from the leaves of the RRIM600 clone located in the clonal gardens of Villavicencio - La Libertad and provided by the Corporación Colombiana de Investigación Agropecuaria (Agrosavia) and. All the samples were obtained under the Addendum No. 20 of the Framework Contract for Access to Genetic Resources and their Derivatives (No. 121 of January 22, 2016) established between the Ministry of Environment and Sustainable Development and the National University of Colombia. The information about each geo-referenced sampling point is shown in Supplementary Table S2.

Propagules were isolated from single foliar, sporulating lesions from which conidia were collected and cultured on M3 solid medium at 25°C in the dark for 45 days until visible stroma formation according to the protocol (82). Once the stroma reached a size of 5×5 mm these were macerated into 2 mL microcentrifuge tubes (Eppendorf®, Germany) and transferred to 125 mL flasks containing M4 sporulation solid medium. The M4 medium consists of potato broth, amino acids, and peptone (82). Sporulation was stimulated by exposing the cultures to white light for 90 minutes for six days (83).

### Genomic DNA extraction and sequencing of *P. ulei*

High molecular weight DNA from sporulated stromata was extracted following Stirling’s protocol (84), modified by adding phenolic extraction followed by three phases of chloroform extractions. DNA concentration, integrity, and purity were assessed by fluorometry (Qubit®), agarose gel electrophoresis (1%) with Tris-borate-EDTA (TBE) buffer stained with SYBR safe (0.5 mg/L), and spectrophotometry (Nanodrop®), respectively. Short-read sequencing was performed using the DNBSEQ Platform Sequencing on the DNBSEQ PE150 instrument by MGI Inc. (China) at the BGI Hong Kong Tech Solution NGS Lab, utilizing the DNBseq DNA library construction kit. Short read raw data was generated in paired-end mode (2×150 bp).

### Genome-wide SNP analyses

To assess genetic variation among *P. ulei* strains, whole-genome resequencing data from six strains collected from different departments in Colombia were analyzed. Raw paired-end reads were quality-checked using FastQC v.0.11.9 (https://www.bioinformatics.babraham.ac.uk/projects/fastqc/), and adapters and low-quality sequences were trimmed using fastp v.1.01 with a minimum quality threshold of Q20 (85). Trimmed reads were aligned to the *P. ulei* reference genome using BWA-MEM v0.7.17 (86). SAM files were converted to BAM and sorted with samtools v.1.20, and duplicates were marked using Picard v.2.27.4 (https://github.com/broadinstitute/picard). Variant calling was performed using GATK (v.4.3.0.0) HaplotypeCaller in GVCF mode for each sample, followed by joint genotyping with GenotypeGVCFs (87). The resulting variant calls were filtered using GATK VariantFiltration based on the following thresholds: QD < 20.0, QUAL < 10000.0, MQ < 30.0, ReadPosRankSum < −2.0 or > 2.0, MQRankSum < −2.0 or > 2.0, and BaseQRankSum < −2.0 or > 2.0. To explore population structure of the six *P. ulei* strains, we performed a principal component analysis (PCA) based on filtered SNPs. The filtered VCF file was first converted to the PLINK binary format using PLINK v.1.9 (88). Linkage disequilibrium pruning was applied with the option --indep-pairwise 50 10 0.2 in PLINK. A PCA was then conducted using the --pca option in PLINK, which generated eigenvalues and eigenvectors. The first two principal components were plotted in R (89), and sample metadata on geographic origin was integrated into the PCA plot to visualize clustering patterns associated with geographic regions.

### Transposable element genome annotation

To obtain high-quality TE libraries for the two largest and most contiguous genomes of species *P. fijiensis* and *P. ulei*, we ran the Earl Grey pipeline v4.1 (90). The Earl Grey pipeline combines identification of TEs based on preexisting libraries and *de novo* approaches for TE annotation. Repetitive elements were first identified and masked by RepeatMasker v4.1.2 (http://www.repeatmasker.org), ignoring low-complexity repeats and small RNA genes. The masked genome was subsequently used for *de novo* TE identification performed with RepeatModeler v.2.0.2 (91) using RepBase v.23.08 and Dfam v.3.3 databases for the DNA and amino acid sequence identification. TEs were classified based on the similarity between *de novo* annotated and known TEs, creating a new combined library. Finally, full-length long terminal repeat retrotransposons (LTRs) were identified with LTR_Finder v1.07 (92). To estimate TE distribution and a potential impact on gene integrity and expression, we compared the annotations of TEs and genes in *P. fijiensis* and *P. ulei* and *Z. tritici* separately. We used BEDtools *closest* v.2.30.0 with the parameter -D a (93). Genes with more than one TE insertions were counted as just a single occurrence.

### Manual TE consensus identification

To obtain high-quality TE family consensus sequences, a manual curation as described in (52) was conducted. In short, the RepeatModeler and Earl Grey consensus sequences were first curated with WICKERsoft (94): similar sequences were searched genome-wide with blastn v.2.13.0 (95). 15-25 sequences of a subset of hits with 300 bp added each up- and downstream were extracted, and a multiple sequence alignment was(G Higgins & M Sharp, 1988)2.1 (96). Visual inspection, as well as information on the sequences of target site duplications and expected start and end sequences were used to define the actual boundaries of each TE family (15), and higher quality consensus sequences were created. New TE families were classified depending on the homology of encoded proteins and the presence and type of terminal repeats, and named after the three letter classification system (15). To remove redundancy and predicted TE families created from TE fragments, each new TE consensus sequence was compared against the already curated consensus sequences with blastn. A large number of previously predicted families turned out to be redundant, as they were fragments of full-length consensus sequences.

A second round TE curation was done to identify non-autonomous TE families that do not contain some or all protein sequences. LArge Retrotransposon Derivates (LARD) and Terminal Repeat retrotransposons In Miniature (TRIM) were detected with LTR-Finder and the filters -d 2001 -D 6000 -l 30 -L 5000 and -d 30 -D 2000 -l 30 -L 500 respectively. Miniature Inverted-repeat Transposable Elements (MITE) were detected with MITE Tracker (97). Short Interspersed Nuclear Elements (SINE) were detected with SINE-Finder in Sine-Scan (98,99). Predicted consensus sequences were compared with WICKERsoft as described above, and removed if less than 5 copies were detected in the whole genome or if a TE consensus sequence already existed. The *P. ulei* reference genome was then annotated with the curated consensus sequences using RepeatMasker with a cut-off value of 250, and simple repeats and low complexity region hits were filtered out.

### Phylogenetic reconstruction of RLG_Mira coding regions

To test if the high-copy TE family RLG_Mira underwent a recent burst, we performed multiple sequence alignment and phylogenetic analyses of its coding regions, following an approach established by Oggenfuss et al. 2023. All full-length sequences and fragments of RLG_Mira copies detected with RepeatMasker in *P. ulei* and a copy from *P. macadamiae* as an outlier were extracted with samtools faidx from the reference genome. Sequences on the negative strand were reverse-complemented. The coding sequence of RLG_Mira was extracted with a blastx search against the PTREP18 TE protein database (https://trep-db.uzh.ch/), and the best hit was retained. A multiple sequence alignment was created containing all sequences from *P. ulei*, the copy from *P. macadamiae* and the coding sequence using MAFFT and the parameters --reorder --local-pair -- maxiterate 1000 -nomemsave--leavegappyregion. The multiple sequence alignment was then trimmed at the start and end positions of the coding sequence using extractalign from EMBOSS. Sequences and fragments that covered less than 50% of the coding region were removed with trimAl v.1.4.rev15 (100). To prevent structural variants in a subset of RLG_Mira copies to from distorting the phylogeny, conserved blocks were extracted with Gblocks v.0.91b, using the parameters -t = d -b3 = 10 -b4 = 5 -b5 = a -b0 = 5 (101). The GC content of each sequence was calculated with geecee in EMBOSS. Maximum likelihood trees were estimated with RAxML v.8.2 (102). First, 10 independent maximum likelihood tree searches were conducted using the parameters with the parameters raxmlHPC-PTHREADS-SSE3 -T 4 -m GTRGAMMA -p 12345 - # 10 --print-identical-sequences. The best maximum likelihood tree was retained. Second, bootstrap analysis was performed to obtain branch support values with the parameters raxmlHPC-P-THREADS-SSE3 -T 4 -m GTRGAMMA -p 12345 -b 12345 -# 50 --print-identical-sequences. Finally, bipartitions were added to the best maximum likelihood tree with the parameters raxmlHPC-PTHREADS-SSE3 -T 4 -m GTRGAMMA -p 12345 -f b --print-identical-sequences. The best scoring maximum likelihood tree was then visualized in R, using read.tree from the package treeio v.1.10.0 to import, ape v.5.7.1 to root the tree based on the *P. macadamiae* copy, tibble v.3.0.1 to add the GC content information to the tree and ggtree(103–106). To detect if RLG_Mira entered *P. ulei* via horizontal transfer, we performed blastx and found that best hits are found in fungi including *Metarhizium anisopliae*.

### Genomic environment of the high-quality reference genome of *P. ulei*

To characterize the genomic environment of *P. ulei*, the largest 12 scaffolds of the reference genome were split into non-overlapping 10 kb windows using EMBOSS splitter v.6.6.0 (107). The percentages coverage by annotated TEs, by the high-copy TE family RLG_Mira and genes per window were calculated using BEDtools intersect v.2.30.0 (93). To calculate a potential impact by RIP mutations, large RIP affected regions in the reference genome were detected using The RIPper (108). The visualization was made with circos (109).

### TE copy number estimation for *P. ulei strains*

The reference genome is not always representative for the whole species, and might be an outlier, which could explain the high TE density. To determine if field strains from different regions contain similar numbers of TEs, we estimated the coverage for each manually curated TE family. Raw reads were first trimmed with Trimmomatic v.0.33 with the parameters: ILLUMINACLIP:TruSeq3-PE-2.fa:2:30:10 LEADING:3 TRAILING:3 SLIDINGWINDOW:4:15 MINLEN:36. Copy numbers for each TE family were then estimated based on normalized coverage and using the method coverage in the McClintock pipeline (110). We attempted to track the positions of the annotated TEs; however, due to their high abundance, it was not possible to identify homologous sites with matches spanning both TEs and non-repetitive genomic regions, preventing their accurate localization within the *P. ulei* genome.

#### Declarations

##### Data availability

Sequence data are deposited at the NCBI Sequence Read Archive under the accession numbers SRR34278610 (*P. ulei*), SRR34278609 (*Pa. egenula*), SRR34278616 (*Rh mozambica*). The additional *P. ulei* isolates were deposited under SRR34278611-SRR34278614 and SRR34278617-SRR34278618. Genome assemblies for *Pa. egenula* and *Rh. mozambica*, gene annotations for the *Pseudocercospora* species and TE annotations for *P. fijiensis* and *P. ulei*, and the TE family consensus sequences are available on Zenodo: https://zenodo.org/records/15862053.

## Acknowledgements

We are grateful to the individuals and institutions who provided natural rubber samples that enabled the isolation of *Pseudocercospora ulei* and the generation of Illumina sequencing data from several departments in Colombia. Specifically, we thank Lyda Constanza Galindo Rodríguez from the University of Amazonia for samples collected in the Caquetá department; the ASOPROCAUCHO Association of Rubber Producers and Traders of Guaviare for samples from the Guaviare department; and Olga María Castro from the Research Group on Conservation Agriculture for Lowland Tropical Soils at AGROSAVIA, La Libertad Research Center, for samples from the Meta department. We thank Tobias Baril from the University of Neuchâtel for help with the Earl Grey pipeline.

## Author contributions

GSSM, UO and DC designed the study. IBA, CAT and FAA provided biological material and performed experiments. AZZ and IS contributed datasets. GSSM and UO conducted analyses. GSSM, UO and DC wrote the manuscript. GSSM and UO acquired funding. UO and DC supervised the work. All authors approved the final manuscript version.

## Supplementary Tables

(see separate file)

Supplementary Table S1: Accession numbers and data resources for the genomes analyzed in this study.

Supplementary Table S2: *Pseudocercospora ulei* isolates analyzed in this study.

## Supplementary Figures

**Supplementary Figure S1.**
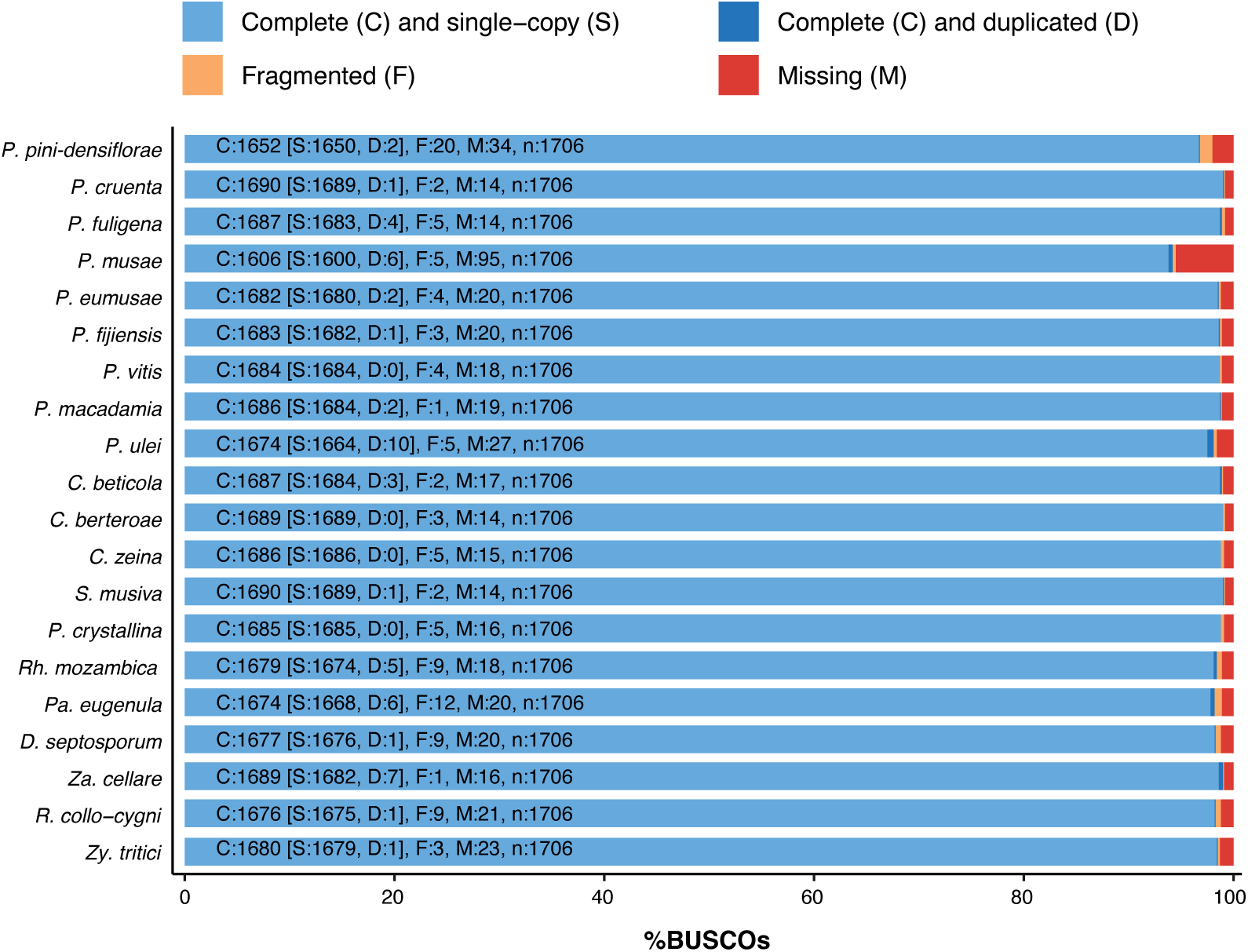
Detailed BUSCO (Benchmarking Universal Single-Copy Orthologs) scores for Mycosphaerellaceae genome assemblies. 1706 BUSCO orthologs from the ascomycota_odb10 database were analyzed, and the complete (single copy or duplicated), fragmented and missing orthologs were listed.

**Supplementary Figure S2.**
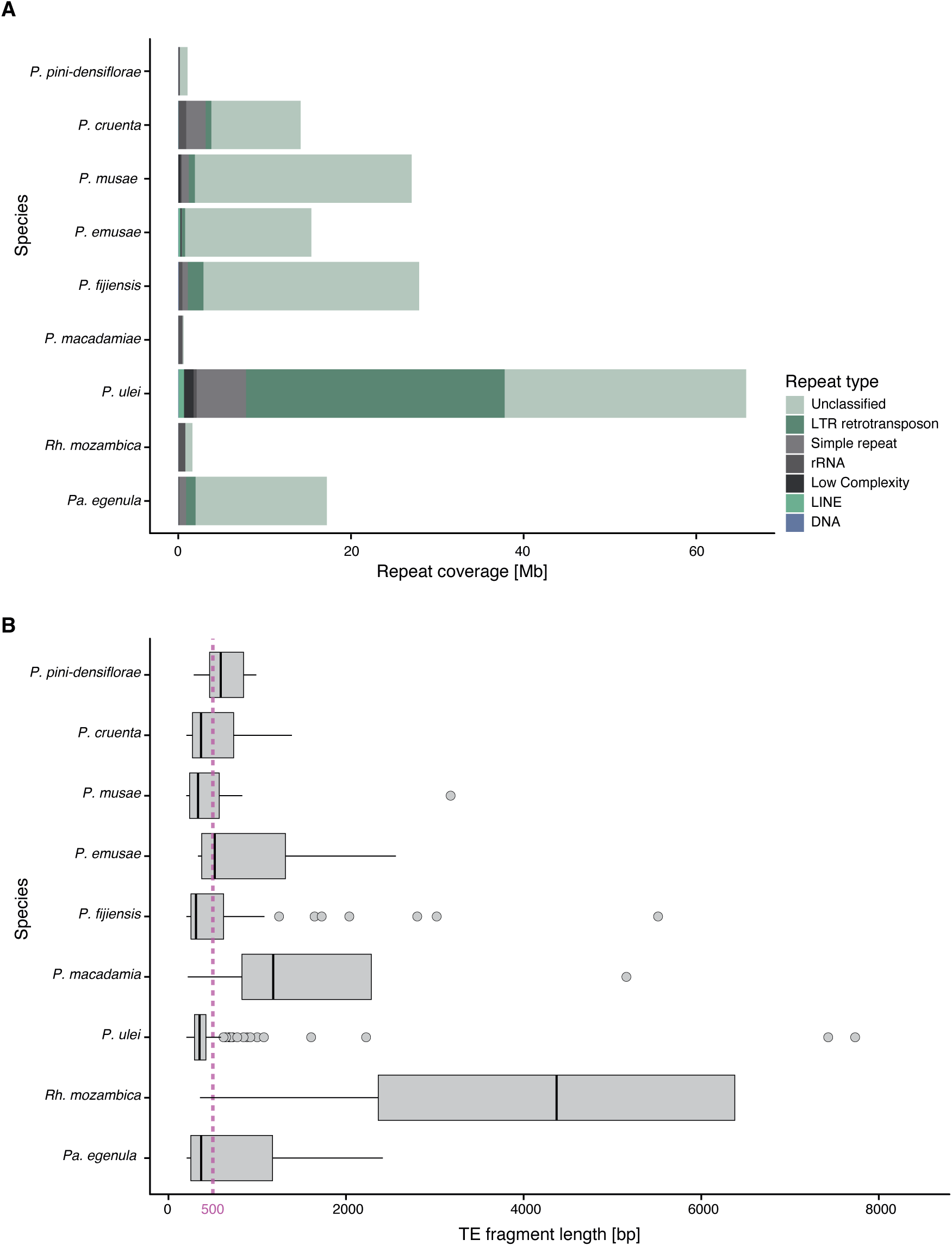
Repeat element lengths in the seven strains with the most expanded genome size of the *Pseudocercospora* genus and two closely related species. A) The total length of repeats in each genome. Colors indicate the type of repeat, with green and blue colors indicating TEs. B) Length distribution of TE fragments per genome. The mauve line indicates 500 bp.

**Supplementary Figure S3.**
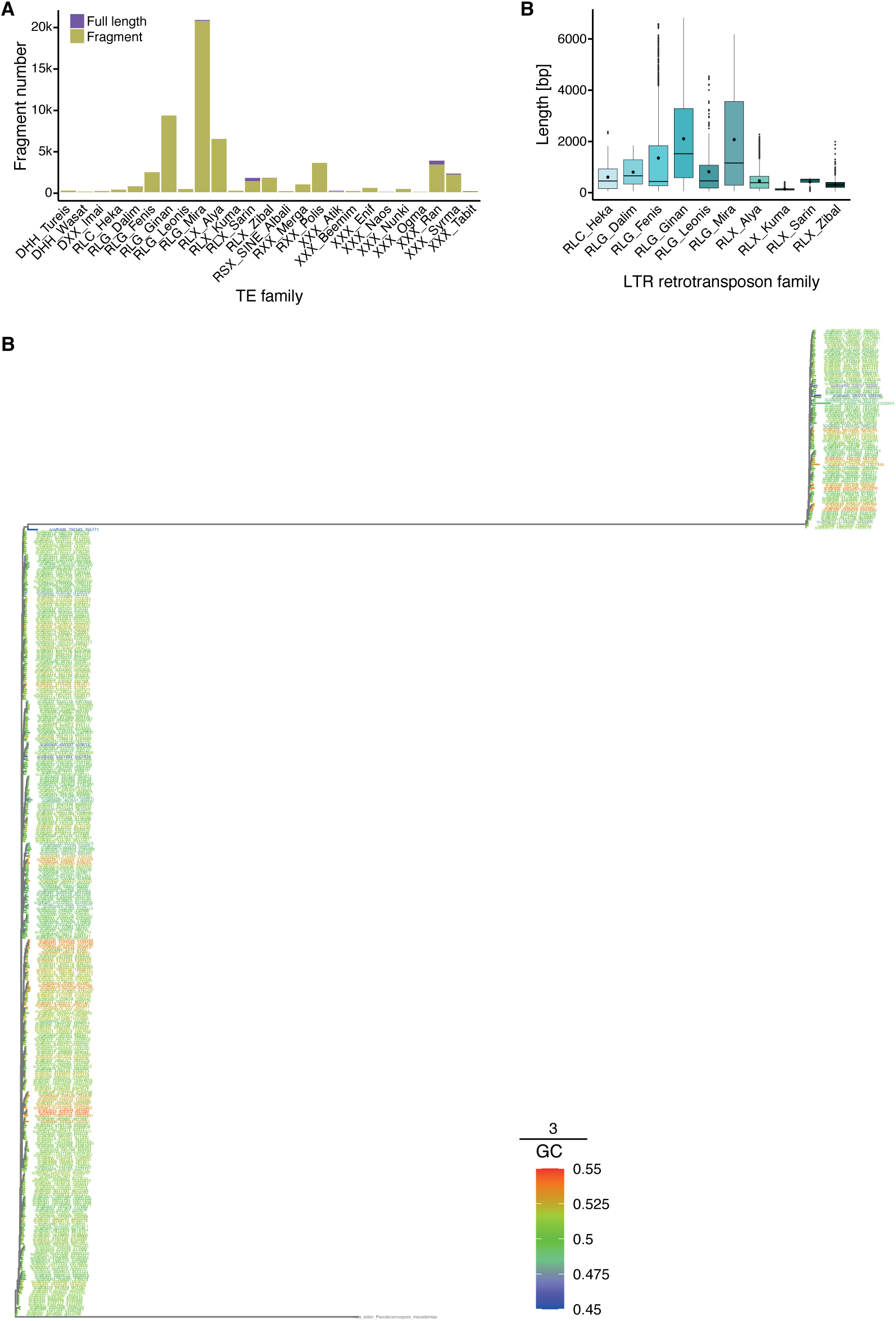
Manually curated TE families in *P. ulei*. A) Copy numbers of TE families. The color indicates if the length of the TE copy was the same (or shorter by >20%) as the consensus. B) Length distribution of TEs for retrotransposons. C) Phylogenetic tree for the coding regions of the RLG_Mira family. The color indicates the GC content, with a low GC content (blue) indicating a potential impact of RIP.

