## Supplementary figures and images for "Massive proliferation of retrotransposons contributes to genome size expansion in species of the *Pseudocercospora* genus"

### Supplementary Figure S1

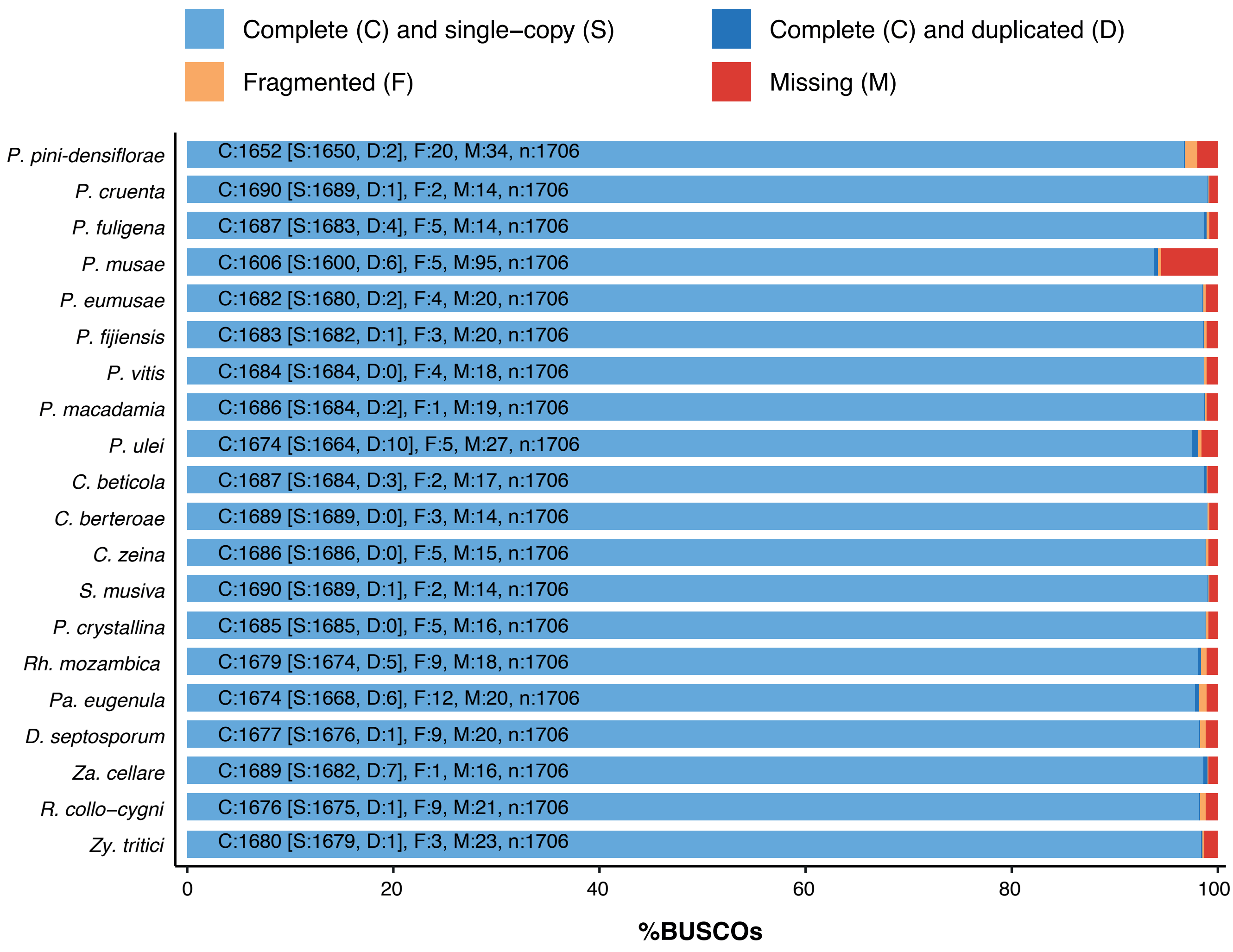

### Supplementary Figure S2

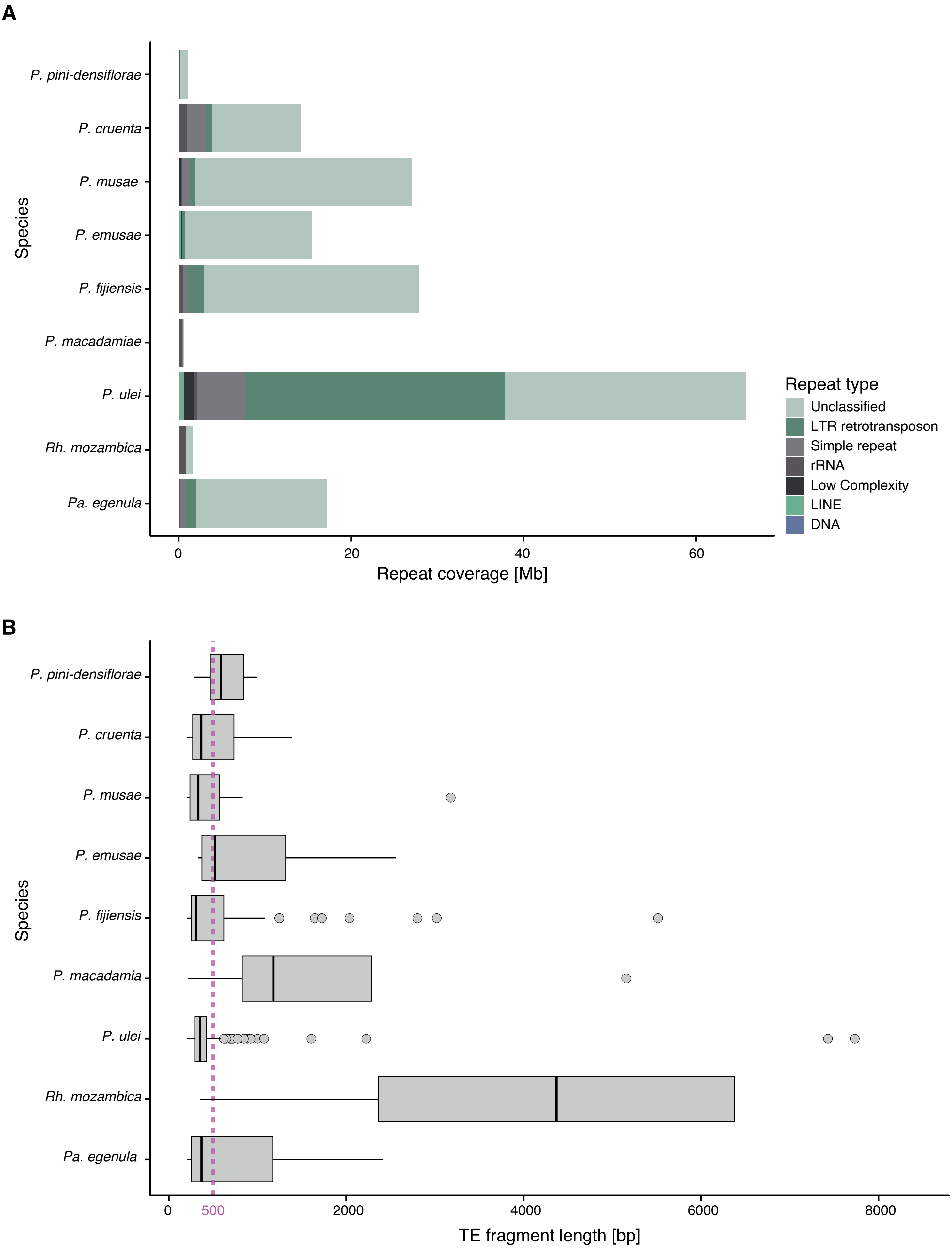

### Supplementary Figure S3

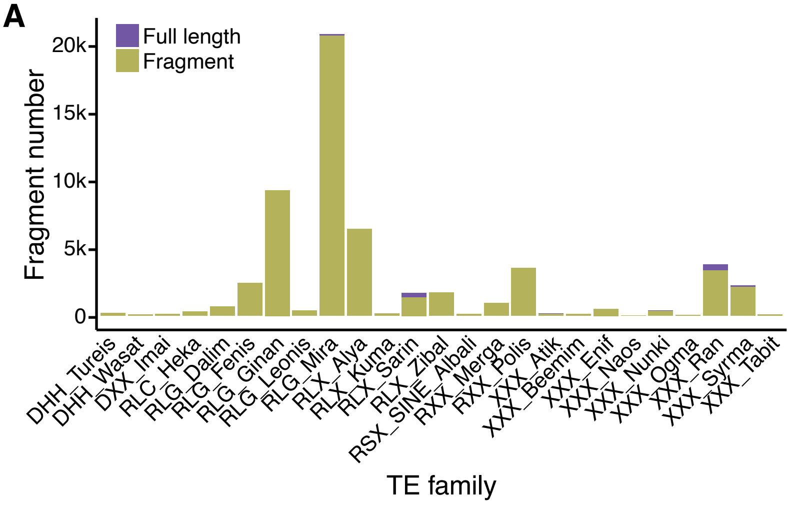

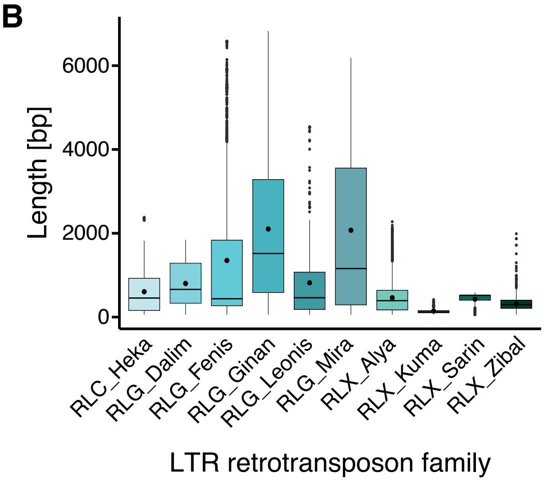

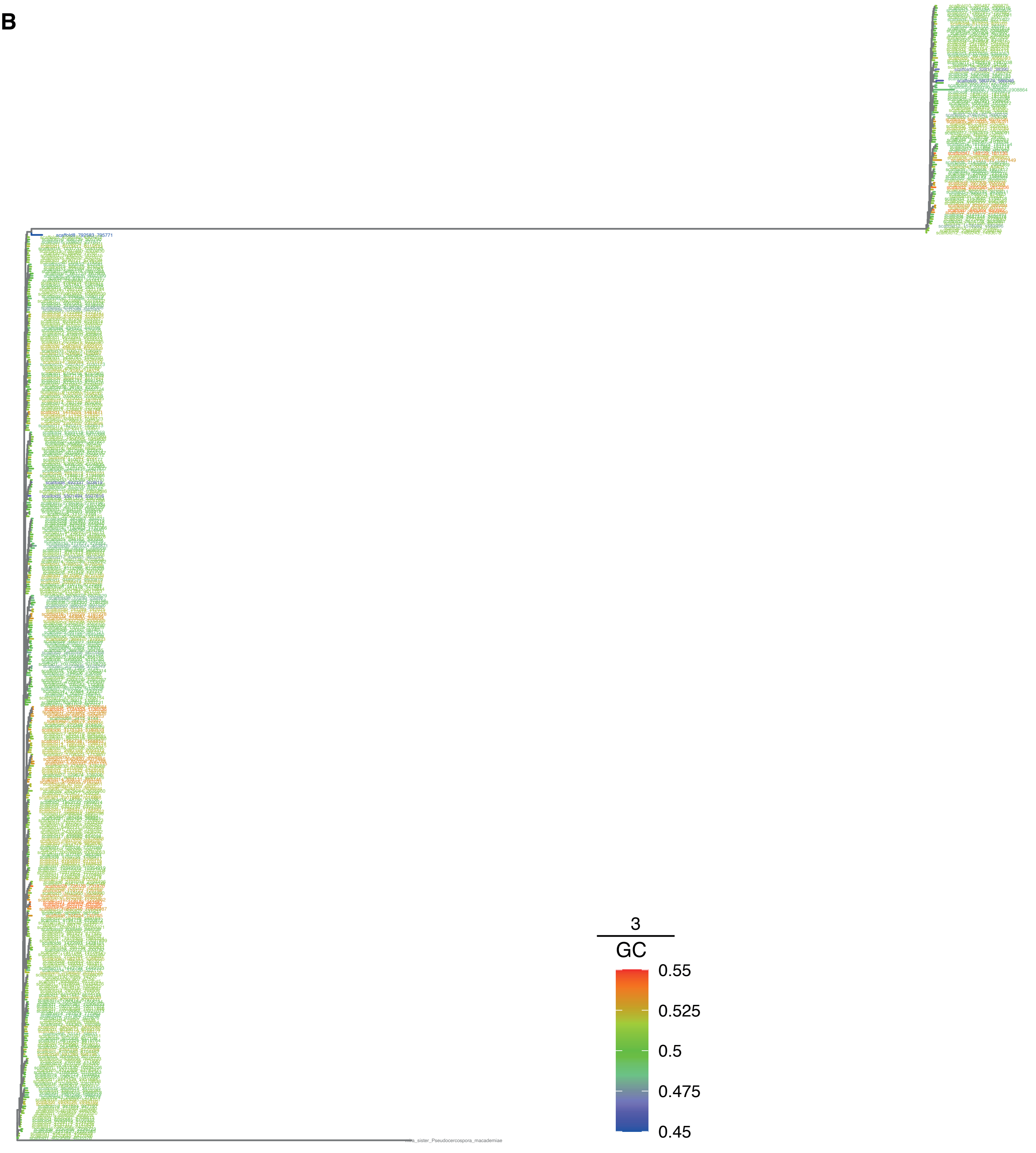
